# A multi-omic cohort as a reference point for promoting a healthy human gut microbiome

**DOI:** 10.1101/585893

**Authors:** Zhuye Jie, Suisha Liang, Qiuxia Ding, Fei Li, Shanmei Tang, Dan Wang, Yuxiang Lin, Peishan Chen, Kaiye Cai, Xuemei Qiu, Qiang Li, Yunli Liao, Dongsheng Zhou, Heng Lian, Yong Zuo, Xiaomin Chen, Weiqiao Rao, Yan Ren, Yuan Wang, Jin Zi, Rong Wang, Hongcheng Zhou, Haorong Lu, Xiaohan Wang, Wei Zhang, Tao Zhang, Liang Xiao, Yang Zong, Weibin Liu, Huanming Yang, Jian Wang, Yong Hou, Xiao Liu, Karsten Kristiansen, Huanzi Zhong, Huijue Jia, Xun Xu

**Affiliations:** BGI-Shenzhen, Shenzhen, China; Shenzhen Key Laboratory of Human Commensal Microorganisms and Health Research, BGI-Shenzhen, Shenzhen, China; Department of Biology, Ole MaalØes Vej 5, University of Copenhagen, Copenhagen, Denmark; China National Genebank, BGI-Shenzhen, Shenzhen 518120, China; Shenzhen Engineering Laboratory of Detection and Intervention of human intestinal microbiome, BGI-Shenzhen, Shenzhen, China.; BGI-Qingdao, BGI-Shenzhen, Qingdao, 266555, China; James D. Watson Institute of Genome Sciences, Hangzhou, China; Macau University of Science and Technology, Avenida Wai long, Taipa, Macau, China

**Author notes:** To whom correspondence should be addressed. X.X.; Z.J.; H.J. These authors contributed equally to this work (Zhuye Jie, Suisha Liang, Qiuxia Ding).

## Abstract

More than a decade of gut microbiome studies have a common goal for human health. As most of the disease studies sample the elderly or the middle-aged, a reference cohort for young individuals has been lacking. It is also not clear what other omics data need to be measured to better understand the gut microbiome. Here we present high-depth metagenomic shotgun sequencing data for the fecal microbiome together with other omics data in a cohort of 2,183 adults, and observe a number of vitamins, hormones, amino acids and trace elements to correlate with the gut microbiome and cluster with T cell receptors. Associations with physical fitness, sleeping habits and dairy consumption are identified in this large multi-omic cohort. Many of the associations are validated in an additional cohort of 1,404 individuals. Our comprehensive data are poised to advise future study designs to better understand and manage our gut microbiome both in population and in mechanistic investigations.

---

The gut microbiome has been implicated in a growing list of complex diseases, showing great potential for the diagnosis and treatment of metabolic, autoimmune and neurological diseases as well as cancer. While case-control studies have been illuminating ^1^, recently published studies have emphasized difficulty in extrapolating to natural cohorts due to heterogeneity in location and ethnicity ^2^’^3^. So far only a few cohorts made use of metagenomic shotgun sequencing instead of 16S rRNA gene amplicon sequencing, the largest being the LifeLines Deep cohort (n=1,135, 32 million reads per sample) from the Netherlands ^4–7^ Fecal or plasma metabolites are more or less included in gut microbiome studies, but the conclusions usually did not go beyond short-chain fatty acids (SCFA), amino acids, vitamin B complex or bile acids. Levels of trace elements such as arsenic have been a health concern (https://www.usgs.gov/mission-areas/water-resources/science/arsenic-and-drinking-water?qt-science_center_objects=0#qt-science_center_objects, https://www.fda.gov/food/metals/arsenic-food-and-dietary-supplements), but are unexplored in the microbiome field. Biological sex is a strong determent for the gut microbiome in mice and livestock ^8–10^. The impact of hormones on the human gut microbiome, or vice versa, remains unclear.

As part of the 4D-SZ (trans-omic, with more time points in future studies) cohort, here we present metagenomic shotgun sequencing data sufficient for high-resolution taxonomic and functional profiling (86.1 ± 23.3 million reads per sample) of the fecal microbiome in a cohort of 2,183 adults, along with questionnaire data, physical fitness tests, facial skin features, plasma metabolome and immune repertoire. Trans-omics analyses in this Han Chinese cohort put into context fecal microbiome disease markers, and uncover previously overlooked measurements such as aldosterone, testosterone, trace elements and vitamin A that influence the gut microbiome, which were validated in an additional cohort of 1,400 individuals. Trends for cardiometabolic diseases and colorectal cancer can be seen, despite the average age of 29.6. This is also to our knowledge the largest cohort with facial skin data and immune repertoire data, which would also be of interest for general health management and disease studies.

A recent study casted doubt over the health benefits of probiotic consumption, concluding that colonization of the strains was highly variable between individuals ^11^. Our large cohort unequivocally showed commercial yogurt strains, especially *Streptococcus thermophilus* and *Bifidobacterium animalis* in feces, and suggested beneficial effects in cardiometabolic health.

## Results

### Comprehensive gut microbiome data together with other omics

Fecal samples were collected during a physical examination, and 2,183 samples (Age, 29.6 ± 5.5, average ± stdev) were subjected to metagenomic shotgun sequencing, yielding 82.95 ± 24.26 million high-quality non-human reads per sample (Supplementary Table 1a), ensuring accurate taxonomic and functional profiling. The reads were mapped to a comprehensive human gut microbiome reference gene catalog containing 9.9 million genes (with a saturating mapping rate of 80.1 ± 4.9 %) and then assigned to 1,507 Metagenomic Species (MGSs) ^12–14^ and 2,981 metagenomic linkage groups (MLGs, Kendall’s tau instead of Pearson’s or Spearman’s correlation between genes) (Qin et al, 2012, Jie et al, 2017), to include both known and unknown microbes. Other omics data, including 104 plasma metabolites (3,980 samples), 634 immune indices (PBMC (Peripheral blood mononuclear cells) V(D)J usage and its shannon diversity, 4,120 samples) from buffy coat, 72 basic medical data (body measurements and routine blood test, 2,715 samples), 49 facial skin imaging indices (2,049 samples), 24 physical fitness data (3,833 samples), 18 entries from psychological questionnaire (2,039 samples), and 56 entries from lifestyle questionnaire (3,820 samples) were collected from the same individuals (Fig. 1, Supplementary Tables 1b-d).

**Fig. 1.**
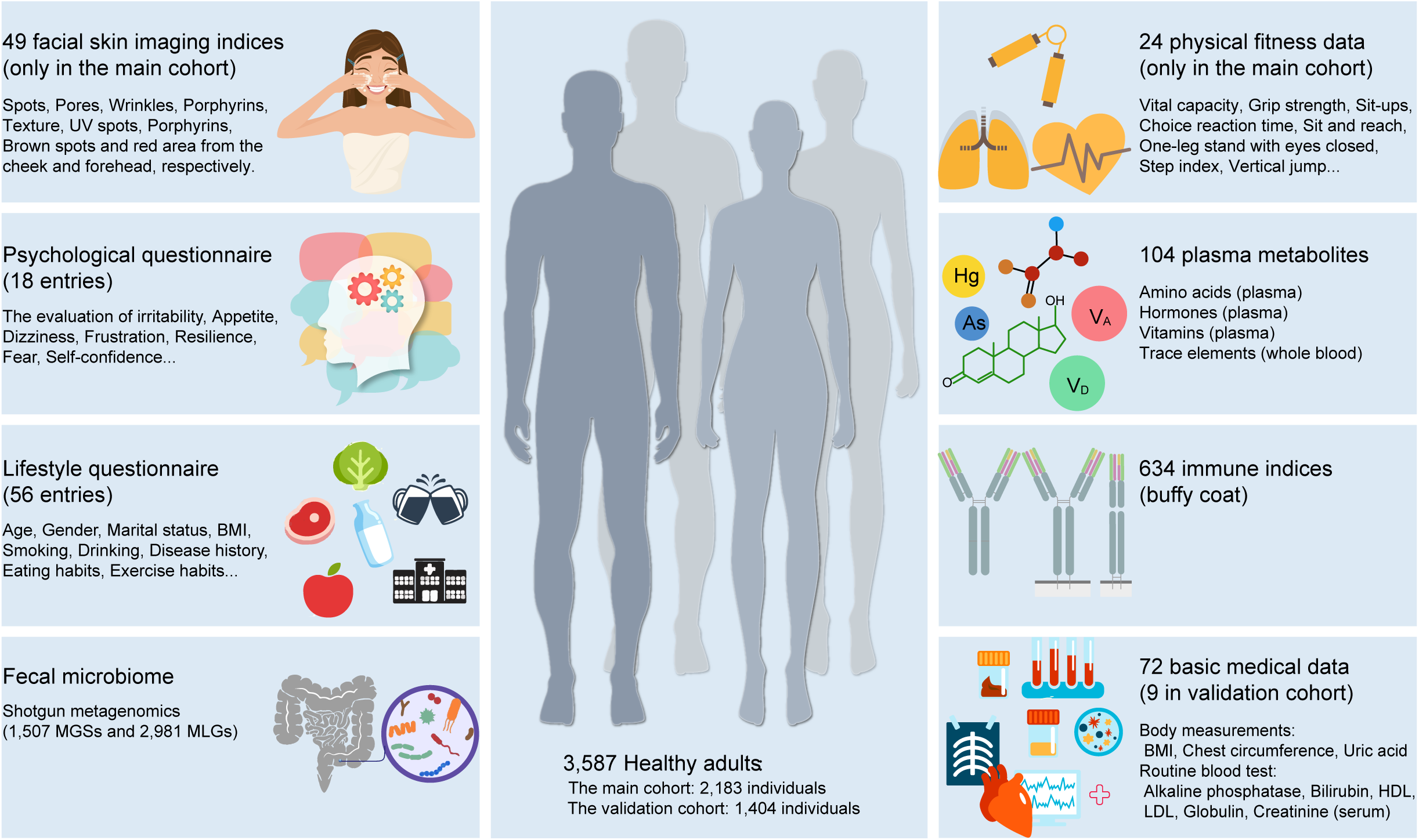
Overview of the multi-omic cohorts. Diagram for features available from the main cohort of 2,183 individuals and validation cohort of 1,404 volunteers. Details are available in Supplementary Table 1.

### The gut microbiome as a relatively independent dimension for health

To get an overall idea of the relationship between omics, an inter-omics prediction value between omics data was calculated using a 5-fold cross-validated random forest model (RFCV, Fig. 2a). Basic medical data showed the highest global systematic association with other omics data. The accuracy of prediction from basic medical data to physical fitness data and from metabolites to basic medical data reaching 75% quantile showed RFCV R = 0.461 and 0.399, respectively (Fig. 2a, b, Supplementary Fig. 1). Basic medical data showed high prediction accuracy to metabolites (Fig. 2a, b); on the other hand, serum creatinine, BMI, waist to hip ratio, hematocrit and triglyceride in basic medical data can be predicted by metabolites (Fig. 2c). Metabolites constituted the highest prediction accuracy to immune indices (R = 0.292) (Fig. 2b). Immune indices showed the second highest prediction accuracy to metabolites (Fig. 2a, b). Facial skin features can be predicted by basic medical data, metabolites, physical fitness data and lifestyle questionnaire (Fig. 2b, c). Among the lifestyle questionnaire, smoking, drinking (especially low concentration alcohol), sports habits (especially resistance training), high-sugar and high-fat dietary habit, and staying up until midnight can be predicted by other omics data (Fig. 2c).

**Fig. 2.**
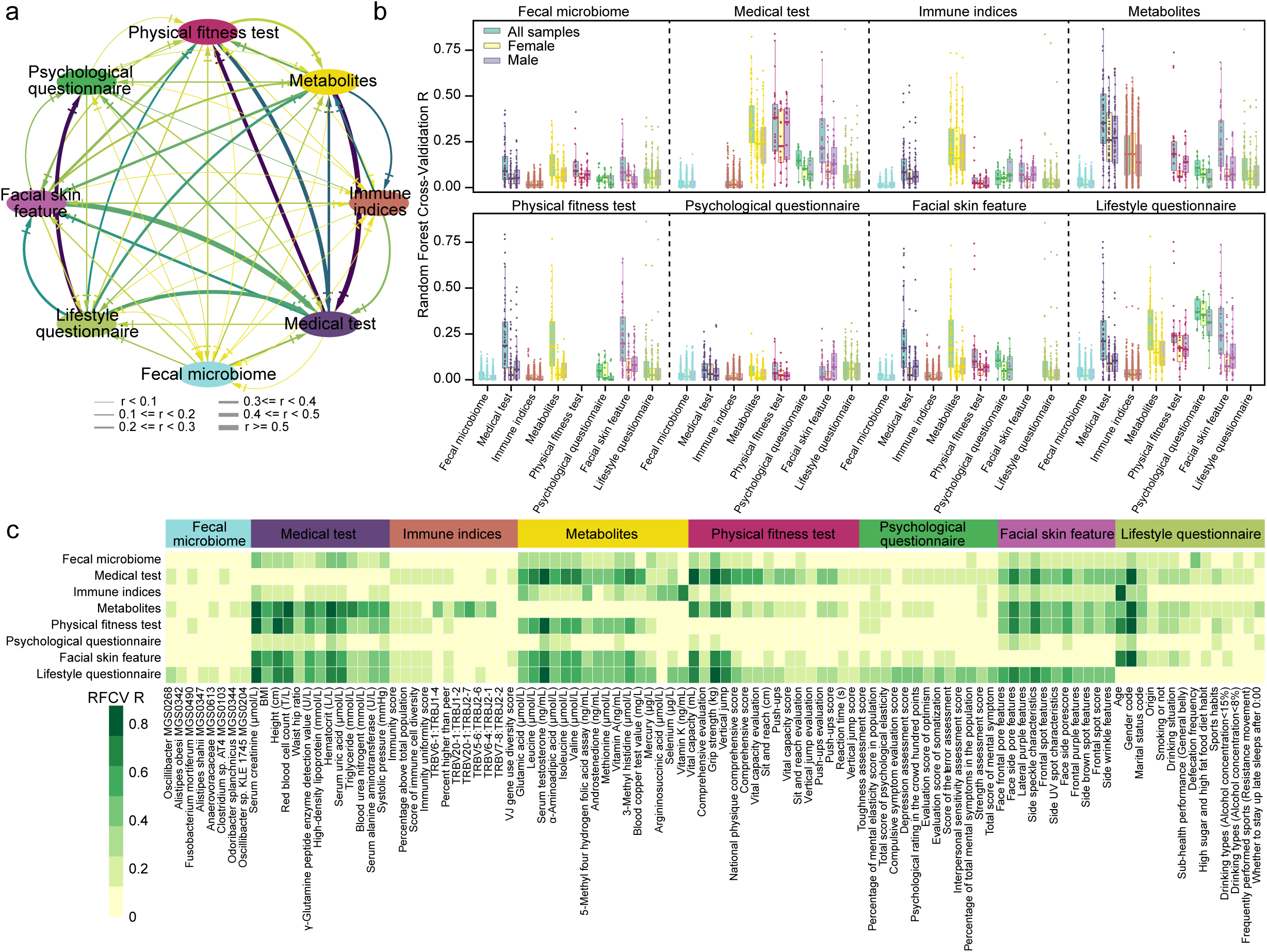
Overview of the interrelationship between omics in the main cohort. **a**, Global association strength between omics datasets. Each arrow is a 5-fold cross-validation random forest (RFCV) prediction. The direction of the arrow indicated the direction of prediction, used the source omics dataset to predict the target dataset. The darkness and size of the arrow lines indicated 75% quantile of spearman’s correlation between measured value and 5-fold cross-validation RFCV predicted value (RFCV R). **b**, Detailed predict power of source omics for each target omics. Tick label in x-axis is target omics. Title in top is source omics. Each node in box is a target factor. The color of the node and box line indicated the target omics data type. Y-axis is the target factor RFCV R predicted from source omics. **c**, Common representative factors that could be strongest predicted by multiple omics data type. Y-axis tick label is source omics. Title is target omics. X-axis tick label is common representative factors (target factors). The cell color in heat map indicated the RFCV R using the omics data in y-axis to predict each factor in x-axis.

A number of factors have been reported to explain gut microbial composition, while the total percentage of variance explained remained in single digits ^4, 15^. According to a RFCV predict model, we observe in this metagenomic cohort influence from lifestyle questionnaire factors such as defecation, yogurt, age, gender, smoking, milk, soymilk, drinking alcohol, fruit and vegetables on gut microbiome composition (Fig. 3), and the cumulative effect size was also in single digits (Supplementary Tables 2b,2c). The BMI distribution is narrow in this cohort (21.729±3.787, Supplementary Table 1b), so its effect size was 0.0015 (q-value=0.014, Supplementary Table 2b). ABO blood group could also predict fecal microbiome composition (RFCV R=0.2, Fig. 3), and specific differences include Lachospiraceae bacterium 3_1_46FAA in blood type A (q = 6.12E-5), *Ruminococcus torques* in blood type B (q = 1.59E-2), unnamed MGS209 in blood type AB (q = 1.59E-2) and *Megaspaera micronuciformis* in blood type O (q = 1.69E-2). As our ‘other genome’, the gut microbiome could predict other omics in this cohort. Gut microbiome showed the greatest prediction power for metabolites, such as plasma vitamins (vitamin A, folic acid, vitamin B5, vitamin D), plasma hormones (testosterone, aldosterone), trace elements (mercury, selenium, arsenic) and plasma amino acids (branched chain amino acids (BCAA), glutamic acid, tryptophan, tyrosine, histidine, alanine) (Fig. 3). Interestingly, hand grip strength, vital capacity, speckles and pores on cheeks and staying up until midnight can also be predicted by the gut microbiome (Fig. 3).

**Fig. 3.**
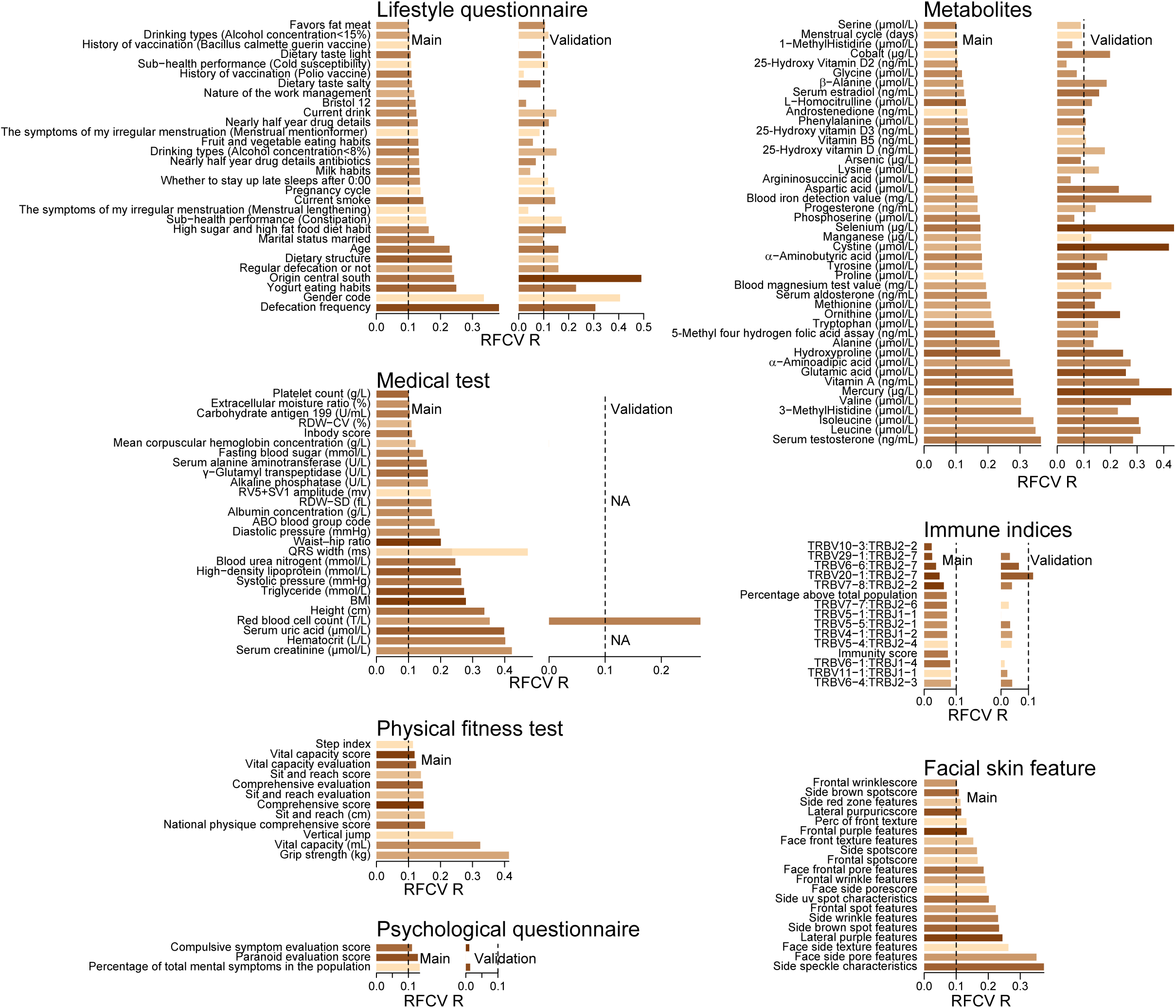
Factors associated with gut microbiome in both cohorts. Top 45 factors with RFCV R > 0.1 in each type of omics that are predicted by gut microbiome. Factors with R ≤ 0.1 in main cohort are not shown. The length of the bar indicated the rank RFCV R using all samples and the color indicated the rank of max of RFCV R using male or female samples only, the darker the greater. Due to missing medical data in the validation cohort (Fig. 1, Supplementary Table 1), only red blood cell count can be validated.

We next included a validation cohort of 1404 individuals (mean age 29.515±5.248, 480 males and 570 females, 82.95 ± 24.26 million high-quality non-human reads per fecal sample), which differed by hometown location compared to the initial cohort (Supplementary Table 1a, Supplementary Table 1b, Fig. 1). The gut microbiome could also predict these plasma metabolites, with greater effects from mercury, cysteine, selenium, iron and cobalt (Fig. 3, Supplementary Table 3), while other data such as physical fitness tests and facial skin features are not available.

### Defecation, hormone and gender

We see that gender (female 1,016, male 1,007) was one of the most significant factors to diverge gut microbiome composition (Supplementary Fig.2a). *Eubacterium dolichum,* and *Blautia wexlerae* were significantly more abundant in males (Supplementary Fig. 2a), after adjusting for age, BMI, medication and dietary supplements (Supplementary Table 3b). *Fusobacterium mortiferum*, which positively associated with testosterone, was sensitive to the statistical adjustments (Supplementary Tables 3a, 3b). Compared to males, females showed a greater α-diversity (Supplementary Table 2a, Supplementary Fig. 2c).

*Bifidobacterium longum*, *B. bifidum*, and *B. catenulatum, B. pseudocatenulatum* were all significantly enriched in females, as well as potentially oral or vaginal bacteria such as *Streptococcus parasanguinis*, *Prevotella bivia* (Supplementary Fig. 2a). Gut microbial functional potential for secondary bile acids strongly associated with self-reported defecation frequency, which were better validated than associations with sex hormones (Supplementary Fig. 2b), suggesting that these are stable patterns.

Aldosterone, one of the major adrenal gland mineralcorticoid, positively correlated with bacteria implicated in cardiometabolic health, such as *Bacteroides intestinalis, B. cellulosilyticus, B. stercorirosoris* and *Eubacterium eligens* (Supplementary Fig. 3) ^16^. *E. eligens* and *Ruminococcus lactaris* scaled negatively with self-reported preference for a salty diet, in contrast to *Blautia obeum* (Supplementary Fig. 2a), and mice on a high salt diet showed decrease in a number of commensal bacteria ^17^.

### The metabolome-immune-gut axis

Among the strongest associations between different omics is that between immune repertoire and plasma metabolites (Fig. 2). More strikingly, when we plotted the associations in detail, the clusters of metabolites corresponded either to the same *TRBV* (T-cell receptor beta variable gene) or to the same *TRBJ* (T-cell receptor beta joining gene) (Supplementary Fig. 4a). Vitamin A, 5-methyl four hydrogen folic acid, selenium, mercury and serum aldosterone showed positive associations with a few TRBJ1-4 and TRBJ2-1, and negative association with TRBJ2-4. Vitamin B5, Vitamin E, phosphoserine, arginosuccinic acid and arsenic showed positive associations with TRBJ1-4, as well as negative associations with TRBJ2-4, TRBV20-1 and TRBV3-1. Glutamic acid and serine showed a pattern that were largely opposite to that of the vitamin A cluster, except for negative associations with TRBV20-1.

We next explored how the gut microbiome might help put the metabolome-immune associations into context. Vitamin A is central to a healthy immune system but is typically studied for its role in early development ^18^. A recent mice study reported modulation of retinol dehydrogenase 7 expression and dampened antimicrobial response in the gut by Clostridiale ^19^. Consistently, we observed associations between Clostridia species (Clostridia MGS0123, MGS0560, MGS0558, Lachnospiraceae bacterium 1_4_56FAA, Lachnospiraceae bacterium 6_1_63FAA, Lachnospiraceae bacterium 9_1_43BFAA, *C, bolteae, Clostridium* sp. AT4, *Clostridium* sp. M62.1) and vitamin A in adult humans both with Spearman’s corelation and with Masaslin associated (Supplementary Fig. 3, Supplementary Table 3a). 5-methyl four hydrogen folic acid exhibited a positive correlation with *Eubacterium eligens* (Supplementary Fig. 4a, Supplementary Fig. 3), a butyrate-producing bacterium that was relatively depleted in atherosclerotic cardiovascular disease ^16^. 5-methyl four hydrogen folic acid also negatively associated with *Dorea* and *Blautia* species (Supplementary Fig. 4a), which have been implicated in obesity and could metabolize formate or hydrogen ^20–22^ (Supplementary Fig. 4a). Associations between the gut microbiome and trace elements including mercury, selenium and arsenic might be surprising (Supplementary Fig. 4a). Selenium-containing rice is commercially promoted as anti-cancer, and we found that the association pattern largely followed arsenic, consistent with these two trace elements’ similar function in anaerobic respiration ^23^. Selenium and mercury also correlated with disease-associated species such as *Clostridium bolteae* and *Ruminococcus gnavus* in the gut microbiome.

The metabolome-immune cluster represented by phosphoserine, and argininosuccinic acidnegatively associated with *Bacteroides coprophilus* (Supplementary Table 3i), a prevalent but not very abundant species from the *Bacteroides* genus. MGSs from *Faecalibacterium prausnitzii* (Supplementary Fig. 4a, Supplementary Fig. 3), a bacterium reported to produce butyrate and metabolize arsenic ^24^, positively associated with L−homocitrulline, phosphoserine, negatively associated with vitamin A, mercury, as well as with specific TCR V(D)J including positive correlation with TRBV27_TRBJ2.3 and TRBV27_TRBJ2.5 and negative correlation with TRBV20−1:TRBJ2−4 (Supplementary Fig. 4a, Supplementary Table 3). The third cluster represented by glutamic acid showed negative associations with previously reported bacteria implicated with lower BMI such as *Alistipes shahii*, *Bacteroides cellulosilyticus*, *Ruminococcus lactis* and *Eubacterium eligens* ^16^ in this large cohort (Supplementary Fig. 4a), consistent with higher glutamic acid in individuals with obesity or insulin resistance ^21, 25^, and here we tentatively identified their associated TCRs (Supplementary Fig. 4a).

Moreover, gut microbiome functional potential showed specific associations with TCR immune repertoire. The gut microbial module (GMM) ^26^ for homoacetogenesis (production of acetate from hydrogen and carbon dioxide) displayed widespread negative associations, most notably with TRBV7-8:TRBJ2-2 (Supplementary Fig. 4b). TRBV7-8 frequency had been reported to be higher in Pima Indian individuals with Type 2 diabetes ^27^ (Supplementary Table 3i). Modules for degradation of arginine and lysine, degradation of lactose and galactose, also associated with a number of VJs (Supplementary Fig. 4b). In the validation cohort, associations with fecal microbiome modules such as lysine degradation, mucin degradation, lactose and galactose degradation, sulfate reduction were validated (Supplementary Fig. 4b, Supplementary Table 3f), which was impressive given the differences in trace metals and other metabolites between the two cohorts (Fig. 3, Supplementary Table 1). So, from both taxonomic and functional points-of-view, the gut microbiome is involved in the metabolome-immune interplay in circulation, with important new leads for experimental investigations.

### Biomarkers for hyperuricemia and cardiometabolic diseases

Hyperuricemia is common in the East Asian population, and urate is excreted in urine or through the gastrointestinal tract. In our cohort, serum uric acid showed negative correlations with gut bacteria such as *Faecalibacterium prausnitzii*, *Alistipes shahii*, *Oscillospiraceae* and *Bacteroides intestinalis* (Fig. 4), adjusted for medication and dietary supplements. Moreover, serum uric acid positively correlated with vitamins (vitamin A, B5, D3 and E), amino acids (glutamic acid and alanine), trace elements (arsenic and mercury), while negatively associated with testosterone (Fig. 4). The negative associations between fecal *Butyricimonas virosa*, *Odoribacter splanchnicus* and plasma alanine were consistent with butyrate production from amino acids (Supplementary Table 3i)^28, 29^, which together with methylhistidines hinted at a meat-excess diet ^30^. Self-reported dietary structure indeed showed association with serum uric acid (Supplementary Table 3j). This is the first set of large-scale evidence for gut microbiome dysbiosis in hyperuricemia, together with hormonal, metabolic and potentially immunological differences.

**Fig. 4.**
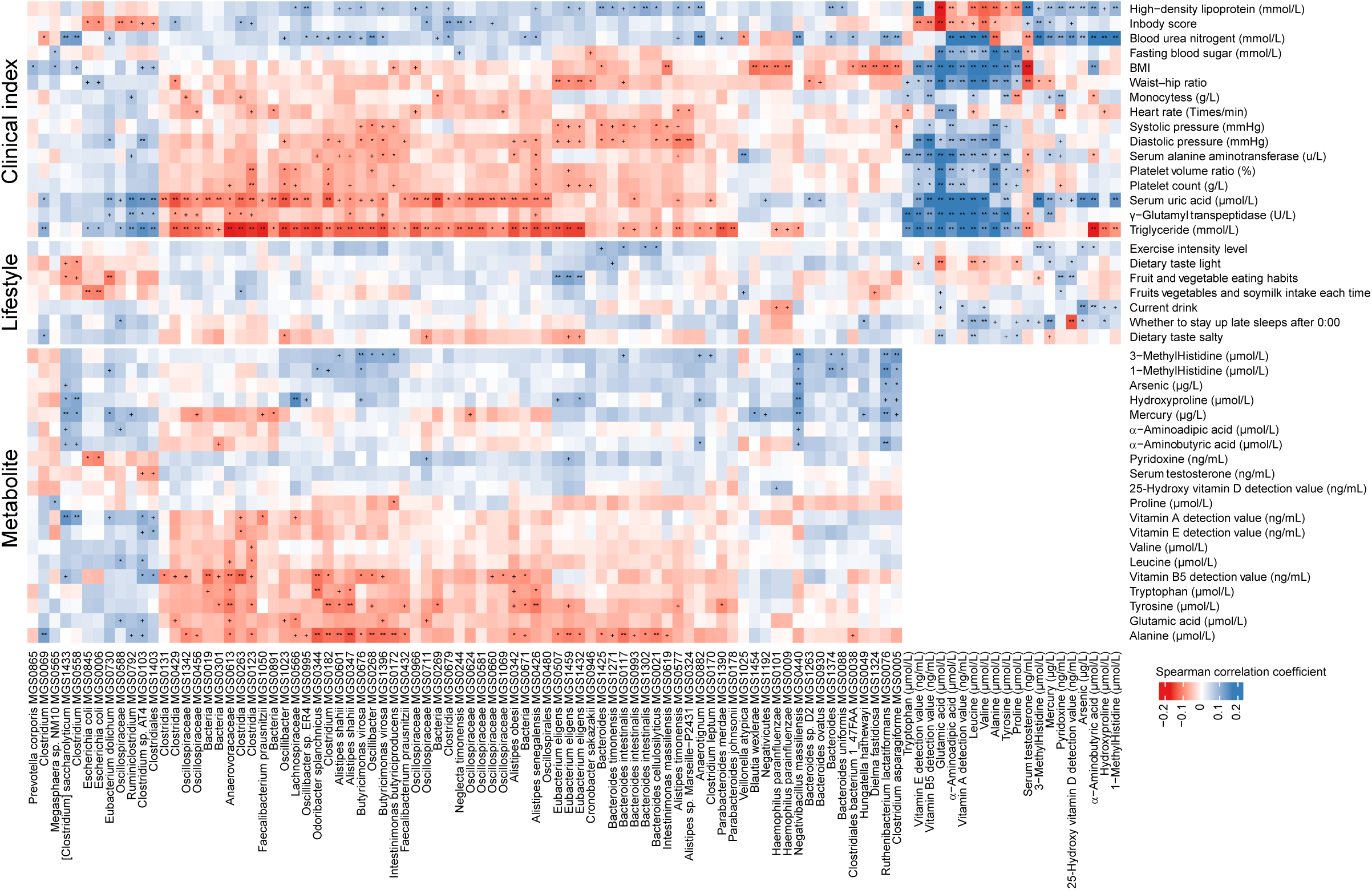
Association map of the four-tiered analyses integrating the metabolites, clinical indices, life style and the fecal microbiome. The color of heat map show the partial spearman correlation adjusted for factors that probably influence the gut microbiome, as shown in Supplementary Fig. 3. BH-adjusted p-value is denoted: +, q-value<0.1; *, q-value<0.05; **, q-value<0.01.

We next defined a score according to 8 routine blood parameters and 80 fecal microbiome features for cardiometabolic disease risk (see Methods) in this young cohort and tested it in previously published case-control samples. With the fecal markers alone, metagenomic samples from Chinese patients with atherosclerotic cardiovascular disease (ACVD), liver cirrhosis, obesity and Crohn’s disease all scored higher compared to control samples without the disease (P <0.05) (Supplementary Fig. 5a), while those from diseases such as colorectal cancer, rheumatoid arthritis and medication-unstratified T2D did not (Supplementary Fig. 5a) ^16, 21, 31–35^. The clinical parameters help clarified T2D and Crohn’s disease (Supplementary Fig. 5b). Thus, although regional differences and misidentifications remain a concern, we illustrate the potential for population-wide screens of cardiometabolic diseases using the fecal microbiome.

### Biomarkers for colorectal cancer

This young multi-omic cohort also provide more insight into the relationship between gut microbiome, plasma metabolome and colorectal cancer (CRC). Both the microbiome and the plasma metabolome are being actively studied for CRC biomarkers, but to our knowledge they have not been investigated in the same cohort. We see here that previously reported CRC-enriched bacteria ^1, 33, 36, 37^ showed associations with plasma metabolites regardless of statistical adjustment for covariates (Supplementary Table 3a). *Peptostreptococcus stomatis* positively associated with plasma leucine, phenylalanine, alanine, tyrosine, as well as sarcosine, a metabolite studied for prostate cancer and a degradation intermediate of glycine betaine ^38, 39^ (Supplementary Fig. 6).

Enterobactericeae including *Escherichia coli, Klebsiella pneumoniae, Enterobacter cloacae* and *Citrobacter freundii* positively associated with sarcosine, hydroxylysine, branched chain amino acids, tyrosine, tryptophan, 1−methylhistidine, hydroxyproline, and argininosuccinic acid (Supplementary Fig. 6). 1-methylhistidine is a marker for habitual meat intake, especially red meat ^30^. Bacteria such as *Bacteroides thetaiotaomicron*, *Butyricimonas virosa* were more associated with 3-methylhistidine (Supplementary Table 3a). Besides, the butyrate-producing *E. eligens* positively associated with fruit and vegetable intake, while negatively associated with plasma alanine (Fig. 4a, Supplementary Fig. 6). A number of these associations were also observed in the validation cohort, e.g. *Enterobacter cloacae* and hydroxylysine, *E. eligens* and alanine (Supplementary Table 3a). These results corroborate fecal markers of CRC with plasma metabolites, and suggest further studies on the long-term interplay between dietary metabolites and bacteria for CRC etiology and threshold for intervention.

### Physical fitness, exercising and sleeping reflected in the gut microbiome

Vital capacity, a commonly used index to assess lung function, positively associated with bacteria such as *A. shahii*, *F. prausnitzii* and *Bifidobacterium adolescentis*, while negatively correlated with disease-related bacteria including *Clostridium clostridioforme*, *Ruminococcus gnavus* and *E. coli*, regardless of statistical adjustments (Fig. 2, Fig. 5, Supplementary Table 3e). Hand grip strength, a protective factor for cardiovascular casualty ^40^, negatively associated with *E. coli* (Fig. 5). Age and sex stratified vertical jump score (Supplementary Table 4) negatively associated with *E. coli*, while positively associated with *B. cellulosilyticus*, *B. intestinalis*, *Eubacterium rectale*, etc. *Bacteroides cellulosilyticus* and *B. stercorirosoris*, which associated with exercise intensity, even correlated with a faster reaction time (Fig. 5), reminding us with associations between *B. cellulosilyticus* and aldosterone, *B. stercorirosoris* and folic acid in both cohorts (Supplementary Table 3a). Moreover, gut microbiome diversity (Shannon index) associated with favorable scores in most of the fitness tests (Supplementary Table 2a).

**Fig. 5.**
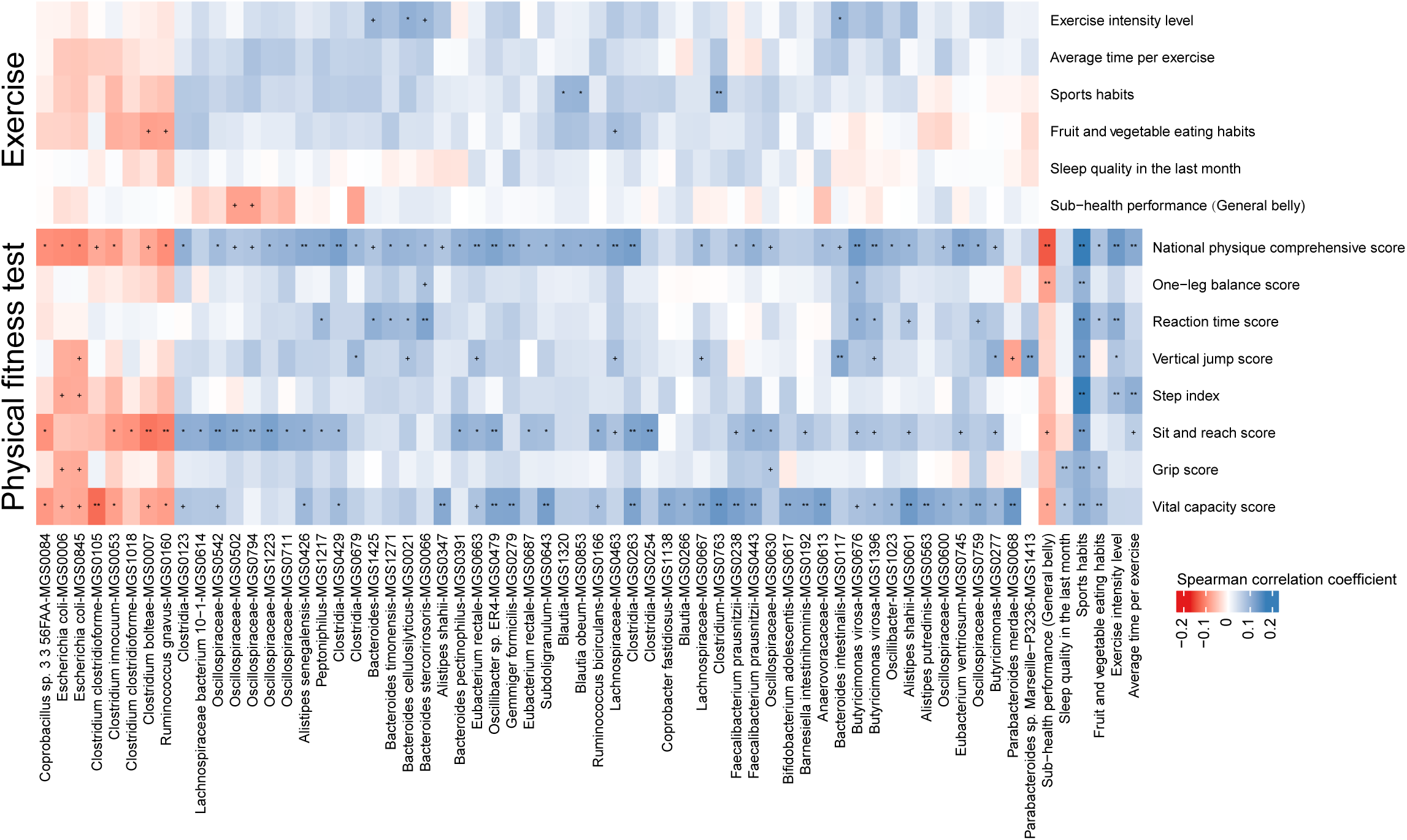
Gut microbiome associated with physical fitness and exercise in the main cohort. The color of heat map shows the partial spearman correlation adjusted factors that probably influence the gut microbiome, as shown in Supplementary Fig. 3. BH adjusted p-value is denoted: +, q-value<0.1; *, q-value<0.05; **, q-value<0.01

Besides, individuals who stay up until after midnight also showed negative correlations with *Holdemania filiformis, Veillonella atypica* and 25-hydroxy vitamin D3/D, while positively correlated with *Clostridium hatheway*, *Clostridium phoceensis*, mercury, selenium, arsenic, vitamin A, hydroxyproline and phosphoserine (Supplementary Fig. 2a, Supplementary Fig. 5, Supplementary Table 3b). Thus, sleeping is also a factor to consider for a complete understanding of the gut microbiome.

### Species from yogurt in the healthy gut microbiome

Besides defecation frequency and gender, yogurt consumption explained a notable portion of variances in the gut microbiome (Fig. 3, Supplementary Table 2b). A recent study casted doubt over the health benefits of probiotics, concluding that colonization of the bacteria was highly variable between individuals ^11^. In both our large cohorts, *Streptococcus thermophilus*, a species included in commercial yogurt mainly for its thermal stability and metabolic support for other strains, was consistently detected in yogurt eaters, and scaled with self-reported frequency of yogurt consumption (Fig. 6, Supplementary Fig. 7). *Bifidobacterium animalis*, likely representing the star strain from CHR HANSEN, *B. animalis* subsp. lactis BB-12, was also enriched in yogurt eaters, and fecal relative abundance of *B. animalis* associated with less stress, less bilirubin, lower diastolic blood pressure, as well as with TCR V(D)J combinations (Fig. 6d), suggesting immune modulation. The association between *B. animalis* and TRBV5.6:TRBJ2.5 was also observed in the validation cohort (Supplementary Table 3c), while the other parameters were unfortunately not available. In contrast to *S. thermophilus*, *B. animalis*, and *Veillonella*, there was no significant increase in any *Lactobacillus* strains (Fig. 6).

**Fig. 6.**
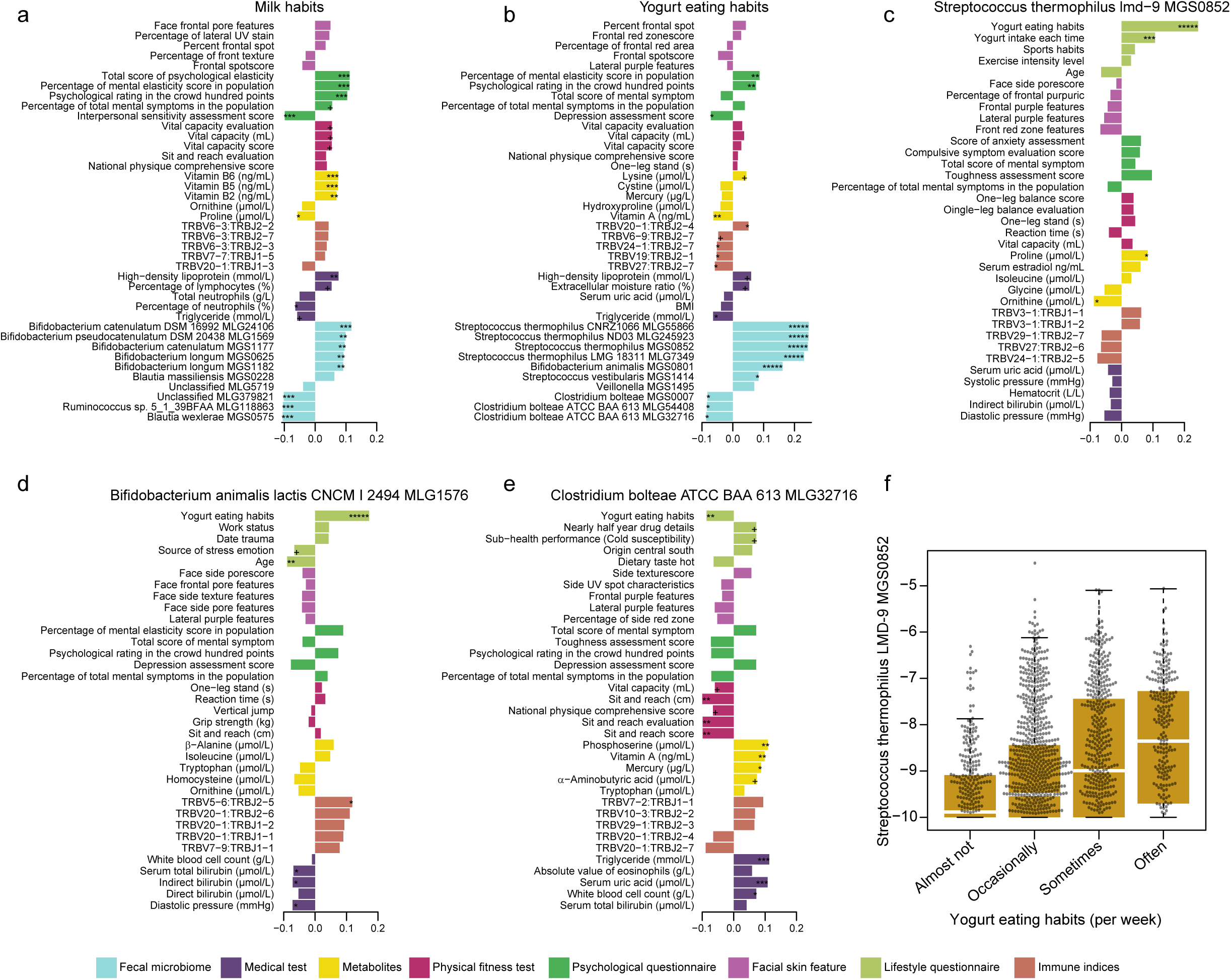
Influence of yogurt and milk intake on omics in the main cohort. **a-e**, The top 5 factor in each omics data associated with yogurt, milk intake habit and the *Streptococcus thermophiles*, *Bifidobacterium animalis and Clostridium bolteae* abundance. The length of the bars represents partial Spearman’s correlation coefficient adjusted for factors that probably influence the gut microbiome, as shown in Supplementary Fig. 3. BH adjusted p-value is denoted: +, q-value<0.1; *, q-value<0.05; **, q-value<0.01; ***, q-value<0.001; ****, q-value<0.0001. **f**, Fecal relative abundance of *S. thermophilus* in volunteers with increasing frequency of yogurt consumption.

Those who used to take yogurt also showed less *Clostridium bolteae*, a bacterium known to be elevated in a number of cardiometabolic diseases ^1, 16^. Intriguingly, fecal *C. bolteae* associated with plasma triglyceride, uric acid, phosphoserine, vitamin A, and mercury (Fig. 6e), offering an explanation for epidemiological evidence of yogurt consumption and reduced risk of gout ^41^. In the validation cohort, *C. bolteae* also associated with mercury and to a lesser extent vitamin A (the vitamin A association was sensitive to covariates, Supplementary Table 3a). Besides, yogurt consumption was associated with a number of favorable measurements such as higher HDL (high-density lipoprotein) cholesterols, lower uric acid and triglycerides, less cysteine, mercury and hydroxyproline (Fig. 6a).

Regarding *Bifidobacterium* in the gut microbiome, however, individuals who consumed milk enriched for *B. longum, B. catenulatum* and *B. pseudocatenulatum* (Fig. 6a, b), implying that some of the yogurt-associated differences come from its exogenous strains such as *S. thermophilus* and *B. animalis*, as well as less *C. bolteae*. The higher *Bifidobacterium* spp., and lower *Blautia wexlerae* and *Ruminococcus* sp. 5_1_39BFAA associated with milk intake were validated in the additional 1404 individuals (Supplementary Table 3). Milk drinking also associated with vitamin B2, B5, B6, HDL, lymphocytes, etc. in the blood, vital capacity, and psychological scores (Fig. 6a, b, Supplementary Fig. 7).

## Discussion

### Insights from multi-omics

In summary, our trans-omic investigation of thousands volunteers establish an unprecedented reference data set for the human gut microbiome. Judging from the associations, it appears as though a number of factors in circulation crosstalk with the gut microbiome, and then manifest on the face, in the head and in fitness tests. Levels of trace elements, such as mercury, arsenic and selenium, as important cofactors for bacteria respiration and other functions ^23^, should be measured even in uncontaminated regions, and in individuals showing normal levels of these elements. Although rice is often studied for such contaminants, exposure can be from other food, drink, air and soil sources^42^(https://www.fda.gov/food/metals/arsenic-food-and-dietary-supplements). Our results suggest that commensal microbial metabolism of trace elements might help determine their levels in the blood, and influence immune functions.

The PBMC TCRβ CDR3 V(D)J usage in such a large cohort is a great resource for discovering microbial antigens other than those from traditional pathogens. While some *TRBV* and *TRBJ* segments are more frequent than others ^43^, we do not yet know how they correspond to T cell sub-populations. Existing studies on TCR have been focusing on pathogens, autoimmune diseases and cancer. For example, TCR profiles of tumor-resident T_reg_ (regulatory T) cells have been shown to significantly overlap with those of circulating T_reg_ cells ^44^; immune phenotype of peripheral blood T_reg_ II cells was not only similar to that of intratumoral T_reg_ cells, but also predicted future relapse of breast cancer patients ^45^. A high diversity in the T cell immune repertoire is believed to be preferable, but the T cell immune repertoire diversity has been reported to be unchanged after a 3-month switch from omnivorous to vegetarian and lower in long-term vegetarians ^46^. In our analyses of this cohort, the overall diversity (Immunity index, Methods) was not the most important factor that predicted other omics, yet could be reflected by metabolites, physical fitness tests, lifestyle and skin features (RFCV R ∼ 0.2, Fig. 2,3). We identify clustering patterns of specific TCRβ CDR3 VJ joining with plasma metabolites including vitamins, trace elements and amino acids (Supplementary Fig. 4a). The chains of causality remain to be fully elucidated; yet, it is likely to be a two-way interplay for metabolite-gut microbiome, metabolite-T cells, and gut microbiome-T cells. Our results imply long-term differences in these features in apparently healthy individuals. A similar speculation could be made for facial skin features, which we expect to be resilient against topical interventions judging from the strong associations with blood parameters.

We have tentatively identified gut bacteria associated with each ABO blood type. A larger proportion of blood type A in Europeans compared to East Asians might help explain the greater abundance of Lachospiraceae bacterium ^12, 47^. Blood type B is more prevalent in northern Chinese, and the blood type B-enriched mucin-degrading bacterium. *R. torques* has recently been reported to show an association with blood glucose ^48^ and was also associated with ulcerative colitis ^49^ and a Bristol stool score of 1 or 2 (Supplementary Fig. 2a). *Megaspaera micronuciformis*, seen in association with blood type O, can produce butyrate from acetate ^50^. Genetic studies of the gut microbiota have not yet reported genome-wide significant associations with ABO blood type genes themselves ^51–54^, while multiple studies have reported impact of *FUT2* secretor/non-secretor status on gut microbiota composition ^55–57^. Tentative associations here are yet to be matched with *in vitro* studies with the glycans ^58, 59^.

Differences in gut microbiome composition between sexes and a greater microbial diversity in females have recently been reported in the LifeLines Deep cohort, yet the gut microbiome in females was influenced by oral contraceptives, ovariectomy as well as antibiotics for vaginal or pelvic infections ^60^. Males of Hadza hunter-gatherers showed differences in gut microbiota compared to females ^61^, including higher *Eubacterium* and *Blautia* in men which were also recapitulated in our Chinese cohort (*E. dolichum, B. wexlerae*). Interestingly, *E. dolichum* associated with a dietary structure of more meat instead of fruit and vegetables, while *B. wexlerae* scaled negatively with milk consumption (Supplementary Fig. 2, Fig. 6). The evolutionary implications remain unclear.

### A baseline of the gut microbiome with deviations towards diseases

Metagenome-wide association studies (MWAS) have documented gut microbial perturbations in a growing list of diseases by comparing cases versus controls. Here we provide a high-depth metagenomic cohort, the mean age for which did not exceed 30 years old. Alarmingly enough, trends for cardiometabolic diseases and colorectal cancer can already been seen from the fecal microbiome and a few parameters in the blood. The set of healthy gut microbes for leanness are increasingly clear ^16^, such as *A. shahii*, *F. prausnitzii*, *E. eligens* and *B. cellulosilyticus*. And we have a better idea how to increase their relative abundances. Interestingly, we observed few association with *Akkermansia*, which may indeed be too diverse among individuals ^5, 62^ or require mucosal sampling. The list of potentially harmful gut microbes are also increasingly clear; future studies are needed to confirm whether we can decrease *E. coli* and *R. gnavus* with exercising and diet, fend off *C. boltae* with yogurt, etc.

While an older cohort would be needed to look at type 2 diabetes ^63, 64^, hyperuricemia is common in this cohort (Supplementary Table 1). *A. shahii* negatively associated with plasma tryptophan (Fig. 4, Supplementary Table 3i), and hyperuricemia has been reported to skew tryptophan metabolism towards kynurenine production in mice models ^65^, instead of indole reported for *A. shahii* ^66^, potentially modulating signaling through aryl hydrocarbon receptors (AhR) ^67^. One of the bacteria negatively associated with serum uric acid, *F. prausnitzii*, has been reported to encode a methyltransferase for arsenic detoxification (Supplementary Table 3i) ^24^. IL-1β, the major cytokine responsible for gout ^68^, has been associated with urinary level of arsenic ^69^. Co-stimulation of patient-derived PBMCs with monosodiurm urate crystals and TLR2 or TLR4 (toll-like receptors) ligands have been shown to disrupt IL-1β/ IL-1Ra (IL1 receptor antagonist) balance ^70^, consistent with involvement of microbes in gout.

Genetic potential for histidine degradation instead of synthesis have been observed to increase in CRC relative to healthy controls according to metagenomic studies ^37, 71^. 1-methylhistidine, a marker for habitual meat intake ^30^, could be metabolized into histidine. Plasma level of the amino acid proline was reported to increase in a mouse model of CRC ^72^, but found in another study to decrease in human CRC ^73^. In this young cohort from China, we did not see significant associations between proline and known gut microbiome markers of CRC. Hydroxyproline, on the other hand, is better predicted by the gut microbiome composition compared to proline (Fig. 3), and associated with meat consumption, staying up until after 0 am (Supplementary Fig. 7). Enterobacteriaceae such as *Escherichia coli* and *Klebsiella pneumoniae* positively associated with hydroxyproline in this cohort. A recent study analyzed fecal metabolites together with fecal microbiome and reported among others an increase in branched chain amino acids and aromatic amino acids in CRC ^74^. Here we observe plasma levels of these amino acids to associate with CRC markers such as *P. stomatis*, and *E. coli*, while the fecal metagenomic potential for leucine biosynthesis was control-enriched ^37, 74^, implying that leucine was normally not in excess.

We also find it intriguing that decarboxylases appear generally important for bacterial stress response in the microbiome, i.e. to maintain a balanced pH for themselves. The top one for gut microbes may be glutamate decarboxylase (produces GABA (γ-aminobutyric acid) from glutamate), while histidine decarboxylase in the female reproductive tract might contribute to menstrual pains ^75^. Besides, recent studies identified tyrosine decarboxylases in gut microbes that could digest the medication levodopa used to treat Parkinson’s disease ^76, 77^.

### Behavioral changes to be trialed for a healthy gut microbiome?

Although effects of sleep fragmentation on hemopoiesis have been seen despite antibiotic treatment ^78^, our results nonetheless suggest that the gut microbiome may have an additional role, together with trace elements, vitamins, and host genetics ^79^. The less hypocretin in mice subjected to sleep fragmentation promoted atherosclerosis ^78^. The increased adiposity and decreased lean mass with sleep loss also involved toll-like receptors (TLRs) ^80, 81^, and we identify cardiometabolic disease-associated species including *Clostridium hatheway* here.

Potential influence of physical activity on the gut microbiota has been analyzed in small cohorts of rugby athletes ^82^ and colorectal cancer ^37^. Although more detailed information for physical activity is preferable, compliance to recordings such as Fitbit is notoriously bad in healthy individuals ^83^. Results from this large cohort at least suggest that exercising might help improve cardio-pulmonary function (grip strength, vital capacity) to decrease incidence of cardiometabolic diseases. Intense exercise, explored for application to individuals with diseases such as prediabetics and Alzheimer’s ^84, 85^, may be no less important than endurance or resistance training; and our results suggest that different types of exercise could have differential impacts on the gut microbiome and the microbiome changes could be a readout for monitoring effects of training. Endurance training actually lowers testosterone ^86^ and could lead to hyperuricemia, especially if combined with high-fructose food and drinks and lack of dairy consumption ^41^.

Our large-scale analyses provide substantial support for health benefits of yogurt consumption. The universally present species were *Streptococcus thermophilus* and *Bifidobacterium animalis* instead of commonly tested probiotics from *Lactobacillus*. An orally administered strain of *B. longum* has been shown to persist in 30% of individuals for at least 6 months ^87^, while we failed to detect in feces an *L. casei* strain gavaged to rats ^88, 89^, suggesting general differences between *Bifidobacterium* and *Lactobacillus*. The strains used by Zmora et al. included a number of *Lactobacillus*, *Bifidobacterium*, as well as *Streptococcus* and *Lactococcus*, all detectable in various gastrointestinal sites despite laxative and colonoscopy ^11^. One potential explanation for the association with desirable cardiometabolic and psychological scores observed in our study for yogurt or milk is the production of metabolites such as folate and GABA by *S. thermophilus*, *Bifidobacterium* and *Lactobacillus* ^90, 91^. Moreover, Lactobacilli have been reported to sequester heavy metals including lead and cadmium ^92^. All of these live or dead probiotics could potentially exert functions on the immune system or even the brain. The positive association with endogenous *Bifidobacterium* species with milk intake is more likely due to live bacteria which help metabolize the lactose in this largely lactose-intolerant population. It remains to be seen whether and how diary consumption affects the gut microbiome in other cohorts, and there appears to be regional differences in China already.

Thus, this study provides a young reference for the gut microbiome with physical fitness test and questionnaire data, and reveals interrelationship with other omics such as trace elements, hormones and immune repertoire that have so far not been included in other study designs. There is a lot more to investigate both in vitro and in vivo by researchers across disciplines. Interventional as well as mechanistic studies will be needed to see how physical activity, well-timed sleeping and dietary interventions such as yogurt, milk and vegetables might improve the gut microbiome, hormone levels, cardiometabolic and mental health.

## Data and materials availability

Metagenomic sequencing data for all samples have been deposited to the CNSA (https://db.cngb.org/cnsa/) of (CNGB) database under the accession code CNP00 00426, CNP0000289.

## Acknowledgments

The authors are very grateful to colleagues at BGI-Shenzhen and China National Genebank (CNGB), Shenzhen for sample collection, DNA extraction, library construction, sequencing, and discussions. We thank Dr. Qiang Sun (University of Toronto), our colleagues Chen Chen and Yanmei Ju, Jinghua Wu for helpful comments.

## Author contributions

J.W. initiated the overall health project. Y.Z., H.Z., K.C., P.C., X.X. organized the sample collection and processing, with immune repertoire from X.L., W.Z, metabolomics from X.Q., Q.L., Y.L., D.Z., H.Lian, Y.Z., X.C., W.R., Y.R., Y.W., J.Z., R.W., raw metagenomic profile from Q.D, X.W., and J.Z., Q.D., S.T., Y.L., D.W. checked the host metadata and matched the omics data. H.Zhou, H.Lu led the DNA extraction and sequencing, respectively. Z.J. led the bioinformatic analyses, including S.L. and F.L. H.Zhong, Q.D., S.T., D.W. performed early analyses which are not included in the manuscript. Z.J., H.J. interpreted the data. H.J., S.L., Z.J., F.L. wrote the manuscript and rendered the display items. All authors contributed to finalizing this manuscript.

## Online Methods

### Study Cohort

As part of 4D-SZ, all the >2000 volunteers for the first cohort were recruited between May 2017 and July 2017 during a physical examination. The 1400 volunteers for the second cohort were also recruited in 2017, with no overlaps. The samples in each omics are shown in Supplementary Table 1c. Baseline characteristics of the cohort are shown in Supplementary Table 1b, 1d.

The study was approved by the Institutional Review Boards (IRB) at BGI-Shenzhen, and all participants provided written informed consent at enrolment.

### Demographic Data Collection

The lifestyle questionnaire contained 56 entries involving age, marital status, disease history of the volunteer and his/her family, eating and exercise habits (Supplementary Table 1b, 1d). The psychological questionnaire contained 18 entries for the evaluation of irritability, dizziness, frustration, fear, appetite, self-confidence, resilience (Supplementary Table 1b).

### Samples Collection

Fecal samples were self-collected by volunteers, using a kit containing a room temperature stabilizing reagent to preserve the metagenome^93^. The samples were frozen on the same day. The overnight fasting blood samples were drawn from a cubital vein of volunteers by the doctors.

### DNA extraction and metagenomics shotgun sequencing

DNA extraction of the stored fecal samples within the next few months was performed as previously described (Qin et al., 2012). Metagenomic sequencing was done on the BGISEQ-500 platform (100bp of singled-end reads for fecal samples and four libraries were constructed for each lane) ^94^.

### Amino Acid Measurements

40 µl plasma was deproteinized with 20 µl 10% (w/v) sulfosalicylic acid (Sigma) containing internal standards, then 120 µl aqueous solution was added. After centrifuged, the supernatant was used for analysis. The analysis was performed by ultra high pressure liquid chromatography (UHPLC) coupled to an AB Sciex Qtrap 5500 mass spectrometry (AB Sciex, US) with the electrospray ionization (ESI) source in positive ion mode. A Waters ACQUITY UPLC HSS T3 column (1.8 µm, 2.1 × 100 mm) was used for amino compound separation with a flow rate at 0.5 ml/min and column temperature of 55 °C. The mobile phases were (A) water containing 0.05% and 0.1% formic acid (v/v), (B) acetonitrile containing 0.05% and 0.1% formic acid (v/v). The gradient elution was 2% B kept for 0.5 min, then changed linearly to 10% B during 1 min, continued up to 35% B in 2 min, increased to 95% B in 0.1 min and maintained for 1.4 min. Multiple Reaction Monitoring (MRM) was used to monitor all amino compounds. The mass parameters were as follows, Curtain gas flow 35 L/min, Collision Gas (CAD) was medium, Ion Source Gas 1 (GS 1) flow 60 l/min, Ion Source Gas 2 (GS 2) flow 60 l/min, IonSpray Voltage (IS) 5500V, temperature 600 °C. All amino compound standards were purchased from sigma and Toronto research chemical (TRC).

### Hormone Measurements

250 µl plasma was diluted with 205 µl aqueous solution, For SPE experiments, HLB (Waters, USA) was activated with 1.0 ml of dichloromethane, acetonitrile, methanol, respectively and was equilibrated with 1.0 ml of water. The pretreated plasma sample was loaded onto the cartridge and was extracted using gravity. Clean up was accomplished by washing the cartridges with 1.0 ml of 25% methanol in water. After drying under vacuum, samples on the cartridges were eluted with 1.0 ml of dichloromethane. The eluted extract was dried under nitrogen and the residual was dissolved with 25% methanol in water and was transferred to an autosampler vial prior to LC–MS/MS analysis. The analysis was performed by UHPLC coupled to an AB Sciex Qtrap 5500 mass spectrometry (AB Sciex, US) with the atmospheric pressure chemical ionization (APCI) source in positive ion mode. A Phenomone Kinetex C18 column (2.6 µm, 2.1 × 50 mm) was used for steroid hormone separation with a flow rate at 0.8 ml/min and column temperature of 55 °C. The mobile phases were (A) water containing 1mM Ammonium acetate, (B) Methanol containing 1mM Ammonium acetate. The gradient elution was 25% B kept for 0.9min, then changed linearly to 40% B during 0.9min, continued up to 70% B in 2 min, increased to 95% B in 0.1 min and maintained for 1.6 min. Multiple Reaction Monitoring (MRM) was used to monitor all steroid hormone compounds. The mass parameters were as follows, Curtain gas flow 35 l/min, Collision Gas (CAD) was medium, Ion Source Gas 1 (GS 1) flow 60 l/min, Ion Source Gas 2 (GS 2) flow 60 l/min, Nebulizer Current (NC) 5, temperature 500 °C. All steroid hormone profiling compound standards were purchased from sigma, Toronto research chemical (TRC), Cerilliant and DR. Ehrenstorfer.

### Trace element Measurements

200 µl of whole blood were transferred into a 15 mL polyethylene tube and diluted 1:25 with a diluent solution consisting of 0.1% (v/v) Triton X-100, 0.1% (v/v) HNO3,2mg/L AU, and internal standards (20 µg/L). The mixture was sonicated for 10min before ICP-MS analysis. Multi-element determination was performed on an Agilent 7700x ICP-MS (Agilent Technologies, Tokyo, Japan) equipped with an octupole reaction system (ORS) collision/reaction cell technology to minimize spectral interferences. The continuous sample introduction system consisted of an autosampler, a quartz torch with a 2.5-mmdiameter injector with a Shield Torch system, a Scott double-pass spray chamber and nickel cones (Agilent Technologies, Tokyo, Japan). A glass concentric MicroMist nebuliser (Agilent Technologies, Tokyo, Japan) was used for the analysis of diluted samples.

### Water-soluble Vitamins Measurements

200 µl plasma were deproteinized with 600 µl methanol (Merck), water, acetic acid (9:1:0.1) containing internal standards, thiamine-(4-methyl-13C-thiazol-5-yl-13C3) hydrochloride (Sigma-Aldrich), levomefolic acid-13C, d3, riboflavin-13C,15N2, 4-pyridoxic acid-d3 and pantothenic acid-13C3,15N hemi calcium salt (Toronto Research Chemicals). 500 µl supernatant were dried by nitrogen flow. 60 µl water were added to the residues, vortexed, the mixture was centrifuged and the supernatant was for analysis. The analysis was performed by UPLC coupled to a Waters Xevo TQ-S Triple Quad mass spectrometry (Waters, USA) with the electrospray ionization (ESI) source in positive ion mode. A Waters ACQUITY UPLC HSS T3 column (1.7 µm, 2.1 × 50 mm) was used for water-soluble vitamins separation with a flow rate at 0.45 ml/min and column temperature of 45 °C. The mobile phases were (A) 0.1 % formic acid in water, (B) 0.1% formic acid in methanol. The following elution gradient was used: 0–1 min,99.0%–99.0% A; 1–1.5 min, 99.0% A–97.0% A; 1.5–2 min, 97.0% A–70.0% A,2–3.5 min, 70% A–30% A; 3.5–4.0 min, 30%A–10.0%A; 4.0–4.8 min, 10%A–10.0%A; 4.9–6.0 min, 99.0%A–99.0%A. Multiple Reaction Monitoring (MRM) was used to monitor all water-soluble vitamins. The mass parameters were as follows, the capillary voltages of 3000V and source temperature of 150°C were adopted. The desolvation temperature was 500°C. The collision gas flow was set at 0.10 ml/min. The cone gas and desolvation gas flow were 150 l/h and 1000 l/h, respectively. All water-soluble vitamins standards were purchased from Sigma-Aldrich (USA).

### Fat-soluble Vitamins Measurements

250 µl plasma were deproteinized with 1000 µl methanol and acetonitrile, (v/v,1:1) (Fisher Chemical) containing internal standards, all-trans-Retinol-d5, 25-HydroxyVitamin-D2-d6, 25-HydroxyVitamin-D3-d6, vitamin K1-d7, α-Tocopherol-d6 (Toronto Research Chemicals). 900 µl supernatant were dried by nitrogen flow. 80 µl 80% acetonitrile were added to the residues, vortexed, the mixture was centrifuged, and the supernatant was used for analysis. The analysis was performed by UPLC coupled to an AB Sciex Qtrap 4500 mass spectrometry (AB Sciex, USA) with the atmospheric pressure chemical ionization (APCI) source in positive ion mode. A Waters ACQUITY UPLC BEH C18 column (1.7 µm, 2.1 × 50 mm) was used for fat-soluble vitamins separation with a flow rate at 0.50 ml/min and column temperature of 45 °C. The mobile phases were (A) 0.1 % formic acid in water, (B) 0.1% formic acid in acetonitrile. The following elution gradient was used: 0–0.5 min,60.0%–60.0% A; 0.5–1.5 min, 60.0% A– 20.0% A; 1.5–2.5 min, 20.0% A–0% A,2.5–4.5 min, 0% A–0% A; 4.5–4.6 min, 0%A–60.0%A; 4.6–5.0 min, 60.0%A–60.0%A. Multiple Reaction Monitoring (MRM) was used to monitor all fat-soluble vitamins. The mass parameters were as follows, Curtain gas flow 30 l/min, Collision Gas (CAD) was medium, Ion Source Gas 1 (GS 1) flow 40 l/min, Ion Source Gas 2 (GS 2) flow 50 l/min, Nebulizer Current (NC) 5, temperature 400 °C. All fat-soluble vitamins standards were purchased from Sigma-Aldrich (USA), Toronto research chemical (TRC).

### Immune indices Measurements

10 ml whole blood was centrifuged at 3,000 r/min for 10 min, then 165 µl buffy coat were obtained to extract DNA using MagPure Buffy Coat DNA Midi KF Kit (Magen, China). The DNA was sequenced on the BGISEQ-500 platform using 200 bp singled-end reads. The data processing was performed using Immune IMonitor ^95^. VJ Gene use diversity is shannon index of VJ gene usage profile. Immune cell diversity is Shannon index of CDR3. Immune cell species result is unique CDR3 number. Immunity uniformity is CDR3 pielou index. Score of above index is the sample rank in population.

### Medical Parameters

All the volunteers were recruited during the physical examination. The medical test including blood tests, urinalysis, routine examination of cervical secretion. All the medical parameters were measured by the physical examination center and shown in Supplementary Table 1b, 1d.

### Facial Skin feature

The volunteer’s frontal face without makeup was photographed by VISIA-CRTM imaging system (Canfield Scientific, Fairfield, NJ, USA) equipped with chin supports and forehead clamps that fix the face during the photographing process and maintain a fixed distance between the volunteers and the camera at all times. Eight indices were obtained including spots, pores, wrinkles, texture, UV spots, porphyrins, brown spots and red area from the cheek and forehead, respectively (Supplementary Table 1b). The percentile of index was calculated based on the index value ranked in the age-matched database, the higher the better (Supplementary Table 1b).

### Physical fitness test

8 kinds of tests were performed to evaluate volunteers’ physical fitness condition (Supplementary Table 1b). Vital capacity was measured by HK6800-FH (Hengkangjiaye, China). Eye-closed and single-legged standing was measured by HK6800-ZL. Choice reaction time was measured by HK6800-FY. Grip strength was measured by HK6800-WL. Sit and reach was measured by HK6800-TQ. Sit-ups was measured by HK6800-YW. Step index was measured by HK6800-TJ. Vertical jump was measured by HK6800-ZT. We got a measure value from each test. Then each measure value score was assigned 1 through 5 based on its corresponding age-matched national standards (Supplementary Table 4). Both the direct measurements and the scores were used for analyses (Supplementary Table 2, Supplementary Table 3).

### Quality control, taxonomic annotation and abundance calculation

The sequencing reads were quality-controlled as described previously ^94^. Taxonomic assignment of the high-quality fecal metagenomic data was performed using the reference gene catalog comprising 9,879,896 gene^12^. Taxonomy of the fecal MGSs/MLGs were then determined from their constituent genes, as previously described^1, 13, 14, 35^.

### The factors in each type of omics predicted by other type omics

The factors in each type of omics were regressed against the relative abundances of mgs profile (found in at least 10% of the samples) in the fecal samples using default parameters in the RFCV function from randomForest package in R. Dichotomous variables (such as gender) and unordered categorical variable (such as region) were re-coding into dummy variables. Frequency items such as yogurt eating habit were assigned integers. RFCV R defined as spearman’s correlation between measured value and 5-fold cross-validation predicted value was calculated, and then rank the top 5 predictable factors in each omics type. The same prediction process was done between any two types of omics. Then ggplot2 package in R was used to boxplot predict power of target omics factors by all kinds of other predictor omics (Fig. 2b). 75% quantile RFCV R between any two types omics (from a to b and from b to a) was used to construct the bi-direction global omics correlation network using CytoScape (Fig. 2a). R pheatmap and barplot was used to make heatmap plot for some representative factors (Fig. 2c, Supplementary Fig. 1).

### Adjusting for potential confounders

Associations between gut microbiome MGSs, functional modules, Shannon diversity, and variance explained and other omics data were all adjusted for factors that probably influence the gut microbiome, including gender, age, BMI, health products (amino acid, vitamin, calcium), antivirus, antibiotics, drugs (currently using antihypertensive drugs, hyperglycemic drugs, lipid lowering drugs), days since last menstrual bleeding, pregnant, lactation, bowel problem (defecation). Besides the above basic set of confounders, we also show the results adjusting for more potential confounders including dietary (dietary taste spicy, sweet, salty, oil, or light, high sugar and high-fat diet habit, fruit and vegetable intake, favors fat meat), exercise (exercise frequency, exercise intensity, average time per exercise), drinking, smoking and Bristol’s stool score.

### Benjamini-Hochberg multiple hypothesis testing correction

The multiple hypothesis testing Benjamini-Hochberg corrections are done for one source target omics pair each time for Fig. 4-6, except immune index and gut microbe pair which BH-adjust was done on one immune index each time. We show two versions of Benjamini-Hochberg correction for Shannon and other omics in Supplementary Table 2a. One of the BH adjust was done within one omics each time. Another adjust was done overall on all omics.

### Robust association network construction between any two omics data type including fecal microbial MGSs

An rank average method ^96^ was used to combine the results of multiple inference methods to make a robust omics association network. We combined two non-linear models, one-to-many randomforest and one-to-one partial spearman’s correlation, to test the association between factor from any two types omics.

*Step 1: Data preprocessing*.

Dichotomous variables (such as gender) and unordered categorical variable (such as region) were re-coding into dummy variables. Frequency items such as yogurt eating habit were assigned integers. We removed variables following these rules: (i) The microbial species less than 10% in all the samples. (ii) Near zero variance. (iii) With more than 70% missing value. Missing values were filled with median. Outliers were defined as outside of the 95% quartiles and outliers samples are removed.

*Step 2: Computation of associations using multiple inference methods*.

For each factor in one omics, we did regression using RFCV function with default parameter based on all factors in one other omics and calculated RFCV R. ^97^. 5-fold average variable importance was output for step3. Partial spearman’s correlation (ppcor R package) between factors from any two types of omics were also output. Potential confounders were considered as described above. We also show generalized linear model results from MaAslin R package^98^) with default parameters after adjusting above confounders.

*Step 3: Robust networks construction*.

To get the robust and strongest association between factors from any two type omics, in other words, to filter predictor factors and target factors, we did it in two steps. First to choose the target factors, we just kept the top 20 target factors with highest RFCV R. Then to choose predictor factors for every selected target factor, we kept predictor factors with top 30 average ranks and retained edges with partial spearman’s correlation BH-adjusted pvalue <0.05. The average rank was computed as sum of the ranks across the RFCV importance and absolute partial spearman rho. For example, metabolites as target and gut microbe as source. We regressed gut microbes against the metabolites and compute the 5-fold cross validation predict power (RFCV R) for each metabolites and partial spearman correlation. 20 metabolites with highest RFCV were kept. For each of the 20 select metabolites such as VA, average ranks across RFCV and partial spearman were done. Gut microbe biomarker for VA was found with average rank top 30^th^ and passed the partial spearman BH-adjusted pvalue <0.05.

*Step 4: Network visualization*.

For each target factor, top 5-10 average ranks source factor in each source omics type were selected as representative factors to make barplots using ggplot2 package (Fig. 6). The pheatmap package was used to plot the common representative factors that could be strongest predicted by multiple omics data type (Fig. 2c). All the source-target factors pair RFCVR (a as source, b as target and b as source, a as target) was boxplot (Fig. 2b) using ggplot2. The ComplexHeatmap package in R was used to plot omics triadic relation (Fig. 4-6). CytoScape was also used to visualize the global omics network (Fig. 2a).

### Microbial metabolic syndrome risk index validation in cardiometabolic cohort

Using multi-omics analyses method described above after controlling for the potential confounders above, we picked up 80 MGSs that significantly correlated with one of the eight cardiometabolic risk factors (waist Hip Ratio, BMI, triglyceride (mmol/L), High-Density-Lipoprotein (mmol/L), serum Uric Acid (μmol/L),γ-glutamyl transpeptidase (U/L), serum alanine aminotransferase(U/L), fasting blood glucose (mmol/L)) (Supplementary Fig. 5, 6). And they are link to the BCAA metabolites (valine / leucine / alanine), tryptophan, glutamic acid (q<0.1, Supplementary Table 3a). For the published disease studies from China, all the MGSs abundances were derived from metagenomic shotgun data, while the 8 clinical measurements could be missing, e.g. liver cirrhosis and Crohn’s disease only had BMI available ^21, 31, 32, 34, 35^ (Supplementary Fig. 5, 6). The microbial metabolic syndrome risk index is similar with the T2D index (Qin et al, 2012). For each individual validation sample, the microbial metabolic syndrome risk index of sample *j* that denoted by *MMSR j* was computed by the formula below:

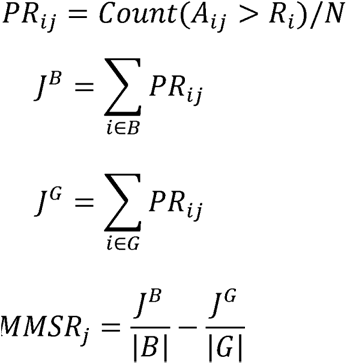

Where *Aij* is a scalar represents the relative abundance of MGS *i* in validation sample *j*.*Ri*; is a vector represents the relative abundance of MGS *i* of all samples in this cohort which served as healthy reference. *N* is the sample size of this cohort that is 2183. Percentile rank PR*ij* is the percentage of test sample *j*’s MGS *i*, relative abundance in its reference cohort frequency distribution that are equal to or lower than it. *B* is 12 out of 80 MGS that were positively correlated with BMI and Triglyceride. *G* is 68 out of 80 MGS that were negatively correlated with BMI and Triglyceride. And |*B*| and |*G*| are the sizes of these two sets. We used percentile rank instead of relative abundance to avoid that the index was influenced too much by the dominant species.

